# *You-Gui-Wan* ameliorates house dust mite-induced allergic asthma via modulating amino acid metabolic disorder and gut dysbiosis

**DOI:** 10.1101/2020.10.15.340646

**Authors:** Wei-Hsiang Hsu, Li-Jen Lin, Yen-Ming Chao, Chung-Kuang Lu, Shung-Te Kao, Yun-Lian Lin

## Abstract

**Introduction:** Allergic asthma is a worldwide health problem, and its etiology remains incompletely understood. Besides, as current therapies for allergic asthma mainly rely on administration of glucocorticoids and have many side effects, new therapy is needed. *You-Gui-Wan* (YGW), a traditional Chinese herbal remedy, has been used for boosting *Yang*, enhancing immunity and treating allergic asthma.

**Objectives:** This study aims to explore the molecular changes during the development of allergic asthma and investigate the potential bio-signatures and the effect of YGW on house dust mite (HDM)-induced chronic allergic asthma in mice.

**Methods:** *Dermatophagoides pteronyssinus* (*Der p*), one of HDMs, was intratracheally administered once a week for a total of 7 treatments over 6 consecutive weeks to induce allergic asthma in mice. Serum metabolomics was analyzed by LC-QTOF-MS/MS. 16S rRNA-based microbiome profiling was used to analyze gut microbiota, and the correlation between metabolomic signatures and microbial community profiling was explored by Spearman correlation analysis.

**Results:** Serum metabolomic analysis revealed that 10 identified metabolites — acetylcarnitine, carnitine, hypoxanthine, tryptophan, phenylalanine, norleucine, isoleucine, betaine, methionine, and valine, were markedly elevated by *Der p*. These metabolites are mainly related to branch-chain amino acid (BCAA) metabolism, aromatic amino acid (AAA) biosynthesis, and phenylalanine metabolism. YGW administration reversed 7 of the 10 identified metabolites and chiefly affected BCAA metabolism. 16S DNA sequencing revealed that YGW profoundly changed *Der p*-induced gut microbiota composition. Multiple correlation analysis indicated 10 selected metabolites have a good correlation with gut microbiota.

**Conclusion:** *Der p* induced BCAA metabolic deviation in allergic asthma mice, and YGW administration effectively ameliorated the AA metabolic disorder, and improved gut dysbiosis. This study paves the way towards the interactions of *Der-p* on microbiome and gut microbiota, and the effects of YGW treatment as well as provides a support for YGW administration with potential benefits for allergic asthma.

## Introduction

Chronic obstructive pulmonary disease (COPD) and chronic asthma represent the worldwide prevalence of long-term airway inflammation. They are among the global public health issues [1]. COPD and chronic asthma result from gene–environment interactions and allergens [2]. One of the major clinical allergens of COPD and chronic asthma is house dust mite (HDM) such as *Dermatophagoides pteronyssinus* (*Der p*) and *D. farina* (*Der f*) [3]. Most patients with COPD and chronic asthma need long-term treatment. The impact on society, economy, and medical expenditure is great and is increasing [4, 5].

Corticosteroids and β-agonists have been used as standard treatment for COPD and chronic asthma. However, they cannot cure these 2 diseases and have side effects, especially in children [5]. An effective treatment with no side effect remains an unmet need. Therefore, many patients search for complementary and alternative therapies for improving health in the remission state and non-acute phase of asthma to prevent acute exacerbations and reduce the dosage of steroids [6].

*You-Gui-Wan* (YGW), an herbal remedy, was firstly mentioned in *Jingyue Quanshu* (1624 A.D.) and has long been used for boosting *Yang* and enhancing immunity in traditional Chinese medicine (TCM) [7]. Previous studies reported that YGW protected the immune function against hydrocortisone-suppressed cytokine expression [8], reduced *Der p*-induced airway hyperresponse and its remodeling, and alleviated allergen-induced airway inflammation in asthma mouse models [9, 10]. However, the action mechanism remains for further investigation.

Because of the complexity of Chinese medicine and the heterogeneity of the etiology of asthma, animals could provide a relative homogenous disease model to explore potential metabolic biomarkers. Metabolomics is a systemic, comprehensive and quantitative analysis of changes in global small-molecule metabolites in a biological matrix, which can be directly coupled to a biological phenotype response to a drug treatment or intervention. It may be a potentially powerful tool to explore the therapeutic basis and clarify the possible action mechanisms of TCM. Therefore, it has attracted much attention in TCM research and is widely used to evaluate the efficacy of TCM in recent years [11].

Gut microbiota associated with metabolic pathways and diseases has been demonstrated [12]. The expression of gut microbial-derived metabolites is associated with the IgE response to allergens and asthma [13]. The gut microbiota composition is related to gut metabolism [12, 14]. Intestinal gut disorders promote the production of metabolic endotoxins, inflammatory factors, and cytokines etc. [15]. Also, diet and TCM could have a beneficial effect on gut dysbiosis and disordered metabolism [16]. These findings highlight the importance of the gut microbiome in disease development.

In this study, we used a comprehensive metabolomics approach to analyze serum and fecal microbiota in mice with *Der p*-induced chronic allergic asthma, the associated metabolic pathway and the association between gut microbiome and metabolomics to explore the potential related mechanisms. We aimed to provide molecular evidence for the beneficial effects of YGW treatment in allergic asthma.

## Materials and Methods

### Chemicals and herbal materials

House dust mite, *Dermatophagoides pteronyssinus* (*Der p*), was purchased from Greer Laboratories (Lenoir, NC). All chemicals were obtained from Sigma (St. Louis, MO) unless otherwise specified.

YGW is composed of Rehmanniae Radix Preparata (root tuber of *Rehmannia glutinosa*), Dioscoreae Rhizoma (rhizome of *Dioscorea opposita*), Corni Fructus (fruit of *Cornus officinalis*), Eucommiae Cortex (bark of *Eucommia ulmoides*), Lycii Fructus (fruit of *Lycium chinense*), Cuscutae Semen (seed of *Cuscuta australis*), Aconiti Lateralis Radix Preparatum (daughter root tuber of *Aconitum carmichaeli*), Cinnamomi Cortex (bark of *Cinnamomum cassia*), Colla Cornus Cervi (antler of *Cervus elaphus*) and Angelicae Sinensis Radix (root of *Angelica sinensis*) at a ratio in the order of 4.4: 2.2: 2.2: 2.2: 2.2: 2.2: 1.1: 1.1: 0.16: 0.16. Each herb was authenticated by Kaiser Pharmaceutics, Tainan, Taiwan and the specimen number as S1 Table. The YGW extract was also supplied by Kaiser Pharmaceutics with good manufacturing practices for pharmaceuticals. The HPLC profile of YGW extract is shown in S1 Fig. The concentrated extract was resuspended in distilled water to produce a final concentration of 100 mg/ml for animal administration [9].

### Statement of animal ethics

6- to 8-Week-old male BALB/c mice (20-22 g) were obtained from the National Laboratory Animal Center in Taiwan. The mice were housed individually and fed a laboratory standard diet (Lab Rodent Chow Die 5001, Ralston Purina Co.) *ad libitum*. Animals were cared and handled according to The Guide for the Care and Use of Laboratory Animals (NIH publication No. 85-12, revised 2010). The animal experiment was reviewed and approved by the Institutional Animal Care and Use Committee of China Medical University (No. 2016-176).

### Animal grouping, drug treatment, and sample collection

BALB/c male mice were randomly divided into 5 groups (n= 4 mice each) for treatment: 1) control (phosphate buffered saline [PBS]); 2) *Der p*; 3) *Der P* + low-dose YGW (0.2 g/kg); 4) *Der p* + high-dose YGW (0.5 g/kg); 5) *Der p* + dexamethasone (Dex, 1 mg/kg). Dex is a synthetic non-selective glucocorticoid (GC) drug that is widely used for immunological, allergic, and inflammatory diseases treatment [17], thence, as a positive control. The dosage of YGW administrated in this study is based on the doctor’s prescriptions which is 2-6 g/70 kg YGW for an adult [18–20].

*Der p*-induced allergic asthma in mice mainly followed a previous method [10]. Briefly, mice were intratracheally administered 40 μl of *Der p* (2.5 μg/μl). *Der p* (in PBS) once a week for a total of 7 treatments over 6 consecutive weeks to induce chronic allergic asthma. YGW and dexamethasone were orally administered daily and once a week, respectively. Both YGW and dexamethasone were orally administered 30 min prior to *Der p* stimulation once a week. After the last treatment, mice were injected with xylazine (200 μg/mouse) and ketamine (2 mg/mouse) in the abdominal cavity and sacrificed [10]. Blood serum samples were collected from brachial artery after anesthesia and stored at −80°C for QTOF-MS analysis. After blood collected, mouse feces were collected from the rectum as soon as possible and kept immediately at −80 ° C.

### Measurement of airway hyperresponsitivity

The airway resistance of mice was measured by using a single-chamber, whole-body plethysmograph (Buxco Electronics, Troy, NY) with doses of methacholine (Sigma-Aldrich, St. Louis, MO) of 0, 3.125, 6.25, 12.5, 25, and 50 mg/ml. Changes in enhanced pause (Penh) represented airway resistance [9].

### Measurement of total IgE and tumor necrosis factor (TNF)-α content

An amount of 100 μl serum was placed in 96-well plates to assess total IgE content with an IgE-specific enzyme-linked immunosorbent assay (ELISA) kit (BD Pharmingen) [10]. TNF-α concentration was determined with a TNF-α ELISA kit (Boster Biological Technology, CA, USA).

### Blood metabolomic profiling

UPLC-QTOF-MS was run on an Agilent 1290 UPLC system (ACQUITY UPLC) coupled with the 6540-Quadrupole-Time-of-Flight (QTOF) mass system (Agilent Technologies, Santa Clara, CA, USA). The untargeted metabolomics profiling was conducted mainly as described [21]. The Trapper package (Institute for Systems Biology) was used to convert MS raw data to an mzXML format. TIPick, an in-house package, was then used to process the mzXML data and remove background signals and detect user-specified metabolites from UHPLC-MS data. Statistical analysis and interpretation focused on only TIPick-identified metabolites. Scaling-based normalization was performed according to the total ion abundance in the UPLC-MS data.

### Bioinformatics analyses

Biological pathway and functional annotation of metabolomics data involved using Protein Analysis Through Evolutionary Relationships Classification System (PANTHER) (http://www.pantherdb.org/), Ingenuity Pathway Analysis (IPA) software (Ingenuity Systems, Mountain View, CA), and MetaboAnalyst 4.0 (www.metaboanalyst.ca/) [22].

### Sequencing of gut microbiota, abundance differences and diversity analyses

Total genomic DNA from fecal samples was extracted by using the column-based method (e.g., QIAamp PowerFecal DNA Kit, Qiagen). DNA quality and quantification were assessed with a Qubit fluorometer (ThermoFisher Scientific) and using the 16S Metagenomic Sequencing Library Preparation protocol from Illumina. Next-generation sequencing involved procedures previously described [23]. Different variable regions of 16S rRNA have been targeted for distinguishing bacteria [24]. The V3-V4 region was identified for distinguishing intestinal bacteria species and was amplified by using a specific primer with a barcode [24]. Reads were quality filtered by using Quantitative Insights Into Microbial Ecology v2 (QIIME2) (v. 2017.10) [25], and chimeric sequences were removed by using UCHIME [26]. The processed sequencing reads (effective tags) were clustered into operational taxonomic units (OTUs) at 97% sequence identity by using UPARSE (http://drive5.com/uparse/) [27]. Then, diversity analysis was performed, and the taxonomy classification of OTUs was assigned according to information retrieved from the Greengenes database [28]. Differences in bacteria abundance were calculated by using linear discriminant analysis (LDA) effect size (LEfSe) [29] and OTU abundance information was normalized by using a standard of sequence number corresponding to the sample with the least sequences. Differences in abundance between groups was tested by the MetaStat method with multiple comparison adjustments [30], and subsequent analysis of alpha and beta diversity involved these output normalized data.

### Statistical analysis

Multivariate statistical analysis of YGW metabolomic data, including principal component analysis (PCA) and partial least-squares discriminant analysis (PLS-DA), was used to analyze the covariance between the measured peak intensities in the MS spectra and the response variables [21]. All analyses involved using IBM SPSS v23.0. Descriptive statistics are presented as mean ± SEM, median (range), or number (%). Student *t*, Fisher exact, or chi-square test was used to compare groups. Paired t test or Wilcoxon signed-rank test was used to compare paired data. All calculated *p*-values were two-tailed. Statistical significance was defined at *p* < 0.05.

## Results

### YGW reduces airway hyperresponsivity in Der p-induced chronic allergic asthma in mice

The allergic asthma model was established by repetitive administration of HDM stimulus to mouse airways as described [9, 10], which resulted in mice with allergic asthma exhibiting airway hyperresponsivity and high total IgE levels, similar to those of clinical symptoms of asthma patients [31]. The disease model group (*Der p*) showed a significant increase in Penh value at both 25 and 50 mg/ml methacholine as compared with the control group (Fig. 1). Moreover, at 25 and 50 mg/ml methacholine, high-dose treatment (*Der p* + YGW (0.5 g/kg)) (oral 0.5 g/kg YGW followed by *Der p* challenge) significantly decreased Penh value as compared with *Der p* alone (Fig. 1A). Furthermore, the positive treatment group (*Der p* + Dex) showed similar effects as the high-dose treatment group (*Der p* + YGW (0.5 g/kg)). Therefore, the high-dose treatment group was used for the following metabolomics and gut microbiota studies.

**Fig. 1.**
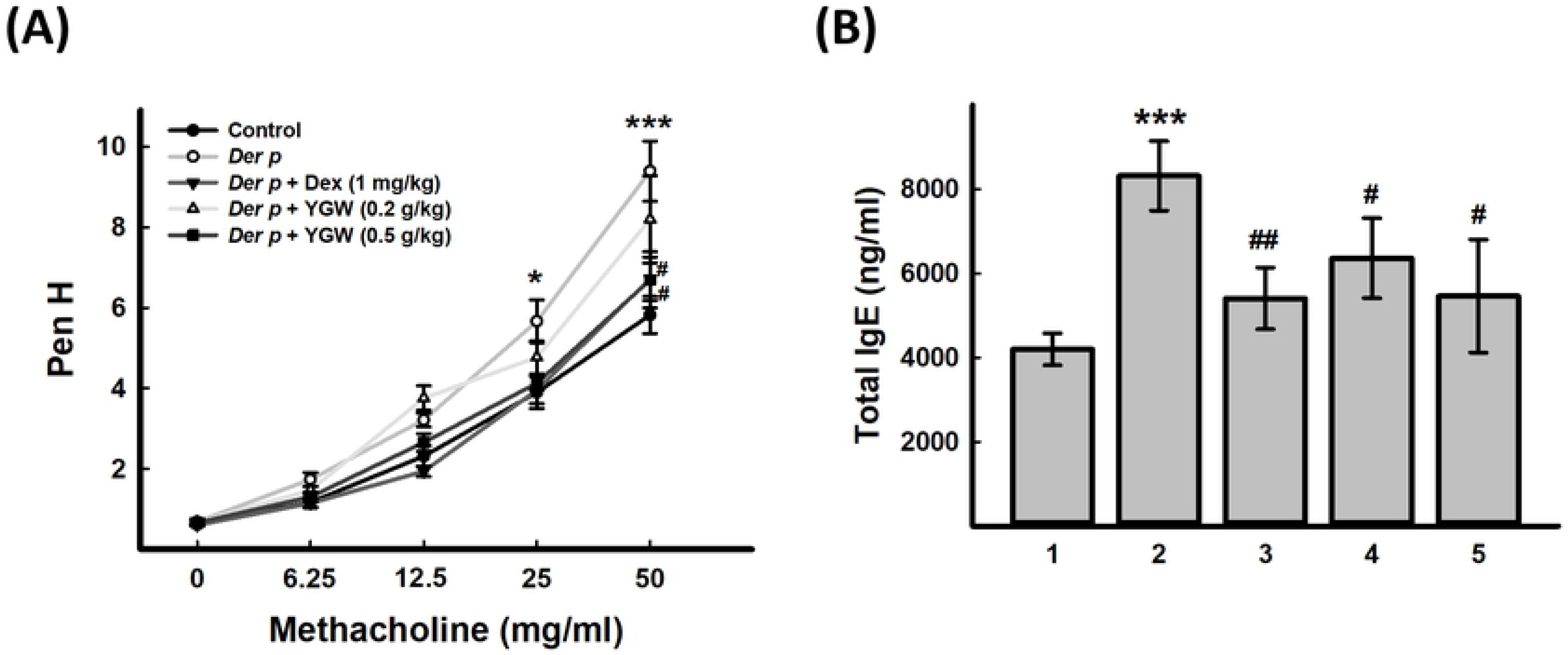
Effect of *You-Gui-Wan* (YGW) on airway hyperresponsiveness and serum total IgE in mice. (A) Evaluation of airway hyperresponsiveness by methylcholine 2 days after last tracheal administration of dust mite, *Dermatophagoides pteronyssinus* (*Der p*). (B) Effect of YGW on total serum IgE level with *Der p*-induced asthma. Data mean ± SEM. *** *p* < 0.001, \**p* < 0.05 (vs control); ##*p* < 0.01, #*P* < 0.05 (vs *Der p*). **1**: control, **2**: *Der p*, **3**: *Der p* + dex (1 mg/kg), **4**: *Der p* + YGW (0.2 g/kg), **5**: *Der p* + YGW (0.5 g/kg)

### Effect of YGW on regulating serum total IgE level with Der p-induced chronic allergic asthma in mice

Total serum IgE content is higher in most patients with asthma than people without asthma [32]. To understand whether YGW regulates humoral immunity, we analysed total IgE content in serum from mice with or without YGW treatment. Total serum IgE content was higher in *Der p*-treated than control mice, and YGW administration could effectively reduce the total IgE content induced by *Der p* (Fig. 1B).

### Effect of YGW on LC-MS-based metabolomics profiles in Der p-induced chronic allergic asthma in mice

To demonstrate the system-wide mechanism of the effect of YGW, we use LC-MS for serum metabolomic analysis. Multivariate statistical methods (PCA and PLS-DA) were used to discern differences among control, *Der p* and treatment groups (*Der p* + YGW (0.5 g/kg)). Exploratory PCA was used to detect intrinsic clustering and possible outliers in the metabolome. The plot of principal component 1 versus 2 scores (S2 Fig. 2A) (R^2^X = 0.900, Q^2^= 0.729) showed an obvious separation among control, *Der p* and treatment groups. By further applying PLS-DA, we observed a reasonably good separation between *Der p* and control groups (R^2^X = 0.896, R^2^Y = 0.661, Q^2^= 0.349) (S2 Fig. 2B). Moreover, findings for the treatment group (*Der p* + YGW (0.5 g/kg)) were closer to the control group, which suggests YGW has a therapeutic effect on the metabolites changed in *Der p*-induced allergic asthma in mice. Also, these results revealed that the models were suitable for predicting the variables that contributed to the class clustering and had low risk of overfitting.

**Fig. 2.**
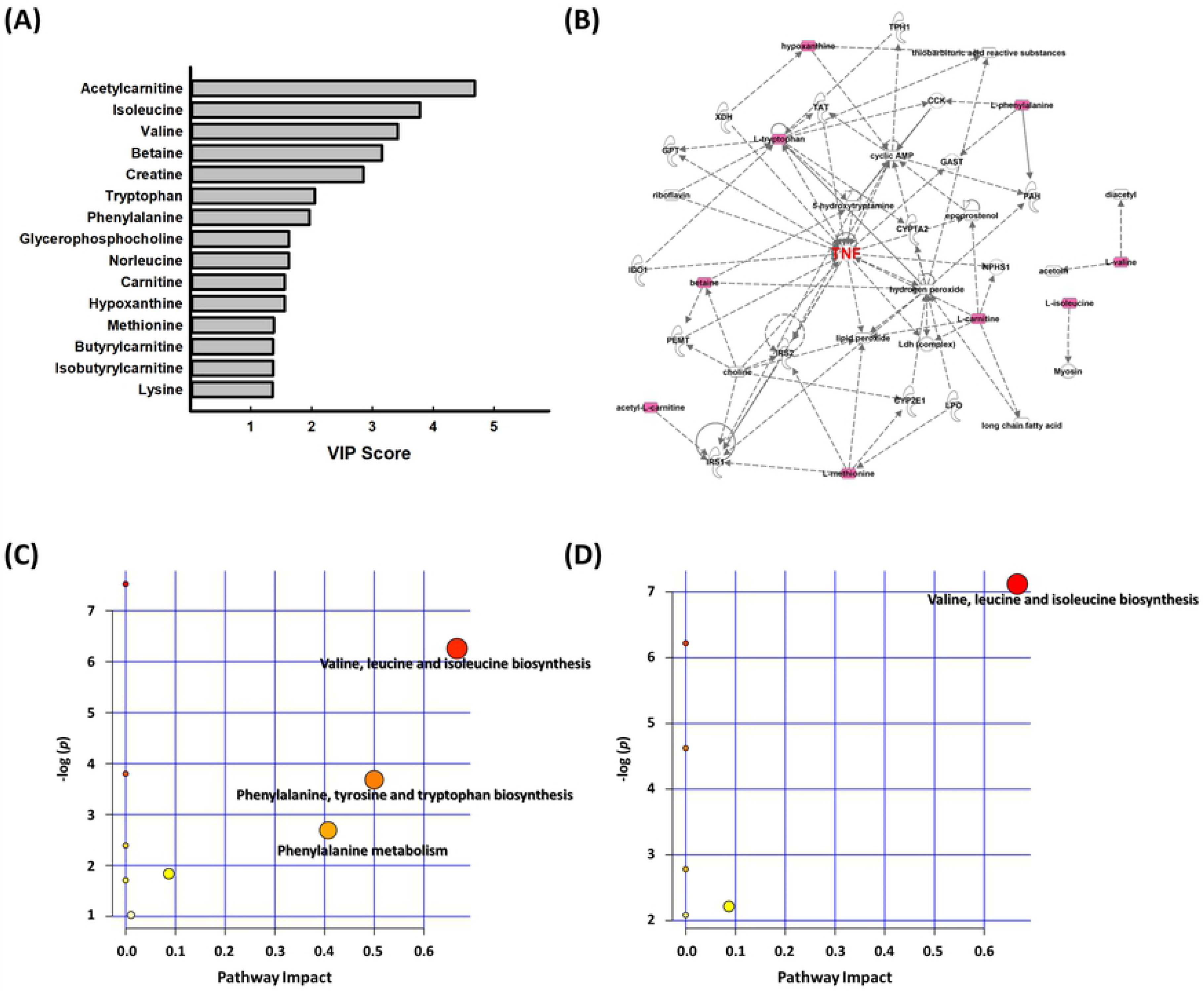
Metabolite signatures from PLS-DA of serum samples in *Der p* and control groups for the variables important projection (VIP) scores and analysis of pathways affected by *Der p*-induced chronic asthma and YGW treatment in mice. (A) VIP analysis based on the weighted coefficients of the PLS-DA model used to rank the contribution of metabolites to the discrimination of *Der p* and control groups by LC-MS/MS. (B) Network pathways identified by using IPA and MetaboAnalyst software. Molecular network of serum of *Der p* group. Direct interactions are represented by continuous lines and indirect interactions by dashed lines. The red nodes represent upregulated metabolites and green nodes downregulated metabolites. Metabolites were inferred in the *Der p* group from changes in serum levels of intermediates during substance metabolism. Pathways affected by (C) *Der p* and (D) high-dose YGW (0.5 g/kg) treatment.

Unpaired *t* test was used to choose metabolites with statistically significant change in expression (*p* < 0.05, control vs *Der p*) (S2 Table). Then, metabolites with VIP score > 1.0 were selected. Significantly *Der p*-altered metabolite signatures included 10 metabolites: acetylcarnitine, carnitine, hypoxanthine, tryptophan, phenylalanine, norleucine, isoleucine, betaine, methionine, and valine (Fig. 2A and Table 1). For 7 of the 10 *Der p*-altered bio-signatures could be significantly reversed by YGW. The level of the remaining 3 metabolites — carnitine, hypoxanthine, and phenylalanine — could also be reversed by YGW but without statistical significance (Table 1).

**Table 1.**
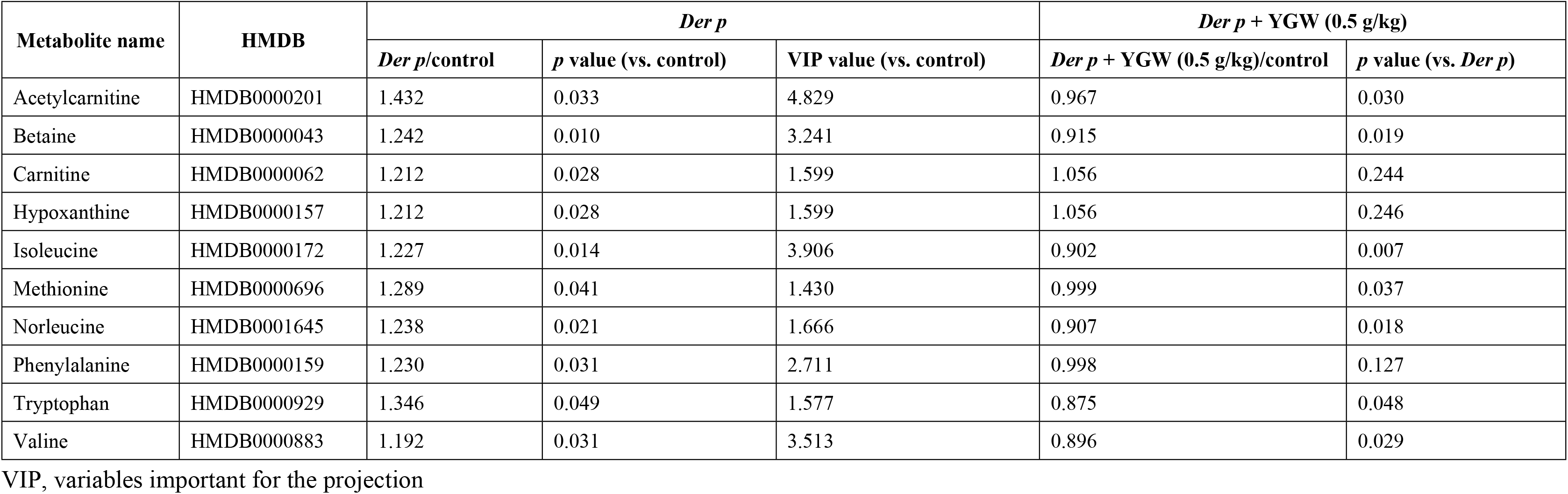
Potential biomarker candidates in *Der p*-altered metabolite signatures.

| Metabolite name | HMDB | *Der p* |  |  | *Der p* + YGW (0.5 g/kg) |  |
| --- | --- | --- | --- | --- | --- | --- |
|  |  | *Der p*/control | *p* value (vs. control) | VIP value (vs. control) | *Der p* + YGW (0.5 g/kg)/control | *p* value (vs. *Der p*) |
| Acetylcarnitine | HMDB0000201 | 1.432 | 0.033 | 4.829 | 0.967 | 0.030 |
| Betaine | HMDB0000043 | 1.242 | 0.010 | 3.241 | 0.915 | 0.019 |
| Carnitine | HMDB0000062 | 1.212 | 0.028 | 1.599 | 1.056 | 0.244 |
| Hypoxanthine | HMDB0000157 | 1.212 | 0.028 | 1.599 | 1.056 | 0.246 |
| Isoleucine | HMDB0000172 | 1.227 | 0.014 | 3.906 | 0.902 | 0.007 |
| Methionine | HMDB0000696 | 1.289 | 0.041 | 1.430 | 0.999 | 0.037 |
| Norleucine | HMDB0001645 | 1.238 | 0.021 | 1.666 | 0.907 | 0.018 |
| Phenylalanine | HMDB0000159 | 1.230 | 0.031 | 2.711 | 0.998 | 0.127 |
| Tryptophan | HMDB0000929 | 1.346 | 0.049 | 1.577 | 0.875 | 0.048 |
| Valine | HMDB0000883 | 1.192 | 0.031 | 3.513 | 0.896 | 0.029 |

VIP, variables important for the projection

### Effect of YGW on metabolic pathway

IPA was used to identify the biochemical pathways which are responsible for the observed metabolic abnormalities. In the network analysis, *Der p*-induced serum metabolites related to allergic asthma tended to gather in a single network. Among the several “hub” molecules at the center of this network, TNF-α was the most important marker (Fig. 2B and S3 Fig.).

To further evaluate the underlying implication of the metabolites with changed expression, we analysed the metabolic pathways by using MetaboAnalyst 4.0 (www.metaboanalyst.ca/). To define the relationships among the metabolites, we generated pathway analysis for the 10 potential biomarkers by using “mouse” as the specific model organism and revealed 7 pathways as most important in *Der p*-induced chronic allergic asthma. 7 metabolites significantly reversed by YGW were used to detect the potential pathway. The most important metabolic pathways (pathway impact > 0.1 and *p* < 0.05) were valine, leucine and isoleucine biosynthesis (branch-chain amino acid [BCAA] metabolism); phenylalanine, tyrosine and tryptophan biosynthesis (aromatic amino acid [AAA] biosynthesis); and phenylalanine metabolism (Fig. 2C). The results suggested that valine, leucine and isoleucine biosynthesis (BCAA metabolism) chiefly contributed to the pharmacological effects of YGW in *Der p*-induced allergic asthma (Fig. 2D).

### Effect of YGW on gut microbiota in Der p-induced mouse chronic allergic asthma

Multiple studies have shown amino acid metabolic disorder is related to gut microbiota [33–35]. To assess the structures changed of fecal microbial communities in control, *Der p,* and high-dose YGW treatment, we generated an average of more than 54,000 sequences from all samples and retained 509 OTUs, then analyzed the diversity of the microbial community (alpha diversity). The average microbiota diversity was slightly lower in *Der p* only than control group, but the low microbiota diversity in *Der p* group was ameliorated by YGW treatment (Fig. 3A).

**Fig. 3.**
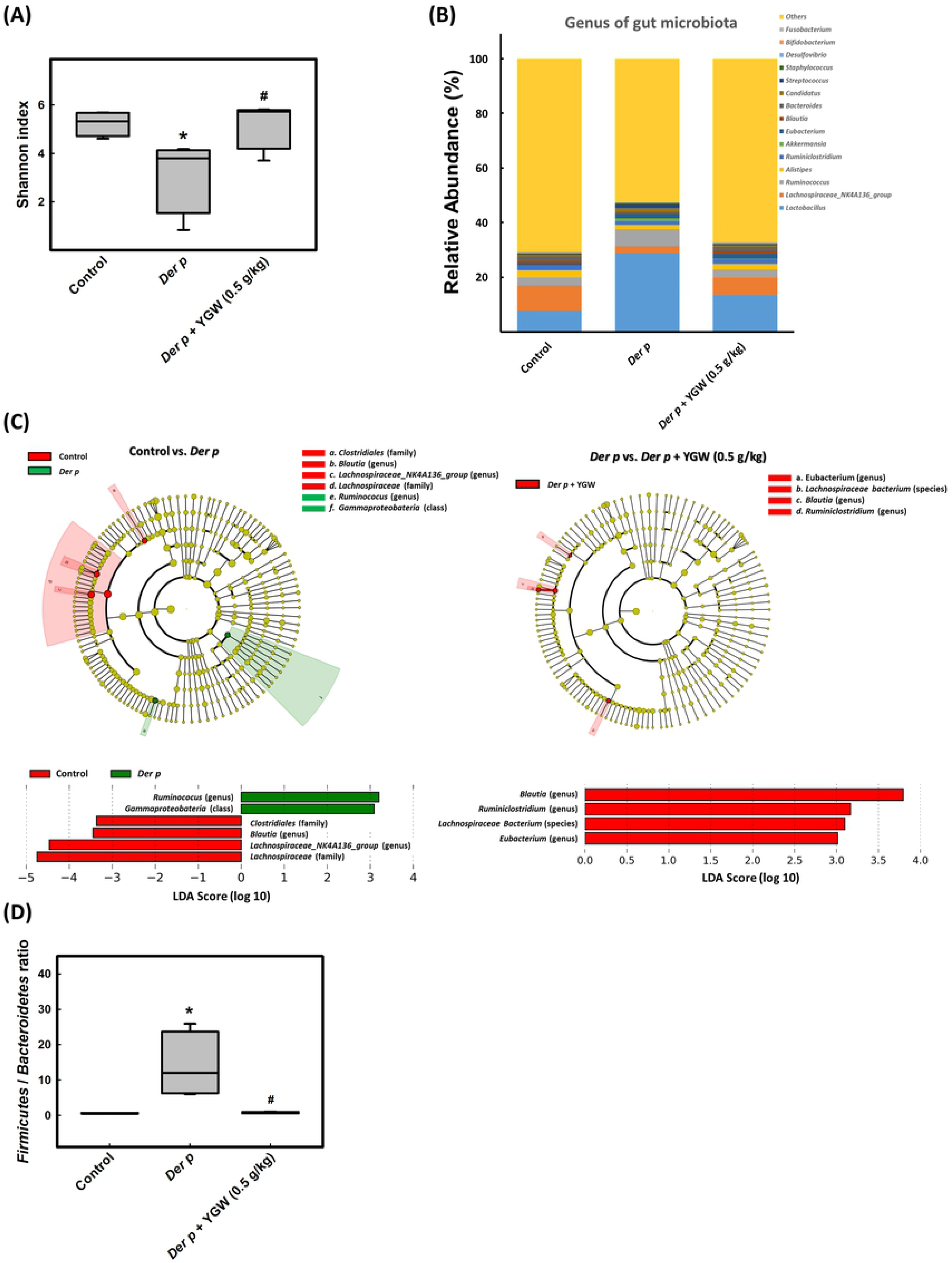
Summary of gut microbiota affected by *Der p*-induced chronic asthma and YGW treatment in mice. (A) Comparison of gut microbiota (Shannon diversity index) among control, *Der p*, and *Der p* + YGW (0.5 g/kg) groups. (B) Gut microbial composition and abundance at the genus level. Each bar represents the top 15 bacterial species ranked by relative abundance. (C) Cladogram generated from LEfSe analysis showing the association between taxon (the levels represent, from inner to outer rings, phylum, class, order, family, and genus). (D) Ratio of *Firmicutes*/*Bacteroidetes* relative abundance. Data are mean ± SEM. \**P* < 0.05 (vs control); #*P* < 0.05 (vs *Der p*)

Furthermore, to intuitively assess the specific changes in the microbial community in the gut microbiota between groups, we analyzed the relative abundance of the dominant taxa identified by sequencing in each group. The top 15 taxa generated the relative abundance superposition histogram at the genus level (Fig. 3B). Among the top 15 taxa, the *Der p* group showed increased relative abundance of *Lactobacillus*, *Ruminococcus*, *Akkermansia*, *Eubacterium*, *Bacteroides*, *Candidatus*, *Streptococcus*, *Staphylococcus*, and *Fusobacterium* but decreased abundance of *Lachnospiraceae_NK4A136_group*, *Alistipes*, *Ruminiclostridium*, *Blautia*, *Desulfovibrio*, and *Bifidobacterium*. YGW treatment partially reversed the change in abundance (Fig. 3B and S4 Fig.). LEfSe analysis, a biomarker discovery tool for high-dimensional data, was used to explore the differences by analysis of taxon abundance in the gut microbiota (from phylum to species) between groups. LDA revealed distinct taxa in the gut microbiota of the groups. A cladogram was generated by LEfSe analysis of sequences from each sample. We identified 9 top differences (LDA score > 3.0) in intestinal flora between control and *Der p* groups and between *Der p* and *Der p*+ YGW groups (Fig. 3C). Our LEfSe analysis revealed that the genera *Clostridiales*, *Blautia*, and *Lachnospiraceae_NK4A136_group* were enriched in control group; *Ruminococus* was enriched in *Der p* group; and *Eubacterium*, *Blautia*, *Ruminiclostridium*, and *Lachnospiraceae_NK4A136_group* were enriched in *Der p*+ YGW group. Therefore, the composition of gut microbiota in different groups was profoundly altered during *Der p*-induced allergic asthma in mice.

Previous study revealed high relative abundance of *Firmicutes/Bacteroidetes* in gut microbiota of asthma patients [36]; thus, we detected whether the ratio of *Firmicutes* to *Bacteroidetes* abundance was increased in *Der p*-induced group. The ratio of *Firmicutes* to *Bacteroidetes* abundance in *Der p*, control, and *Der p*+ YGW groups was 0.59, 11.49, and 0.70, respectively (Fig. 3D), with no significant difference between control and *Der p*+ YGW groups.

### Correlation between metabolomic signatures and microbial community

To investigate the interrelations between the change in metabolite levels and microbiota composition, we used Spearman rank correlation analysis of genera with the highest expression in gut microbiota and the selected serum metabolites, then represented these in a heat map (Fig. 4). Multiple correlation analysis showed a positive correlation (*p* < 0.05) between *Candidatus* and carnitine (r=0.636), hypoxanthine (r=0.636), norleucine (r=0.587), methionine (r=0.643), and tryptophan (r=0.601). However, *Blautia* and *Lachnospiraceae_NK4A136_group* showed a negative correlation (*p* < 0.05) with most of the 10 selected metabolites: acetylcarnitine, carnitine, hypoxanthine, phenylalanine, norleucine, isoleucine, betaine, methionine, and valine. Therefore, the expression of the 10 selected metabolites was well correlated with gut microbiota composition.

**Fig. 4.**
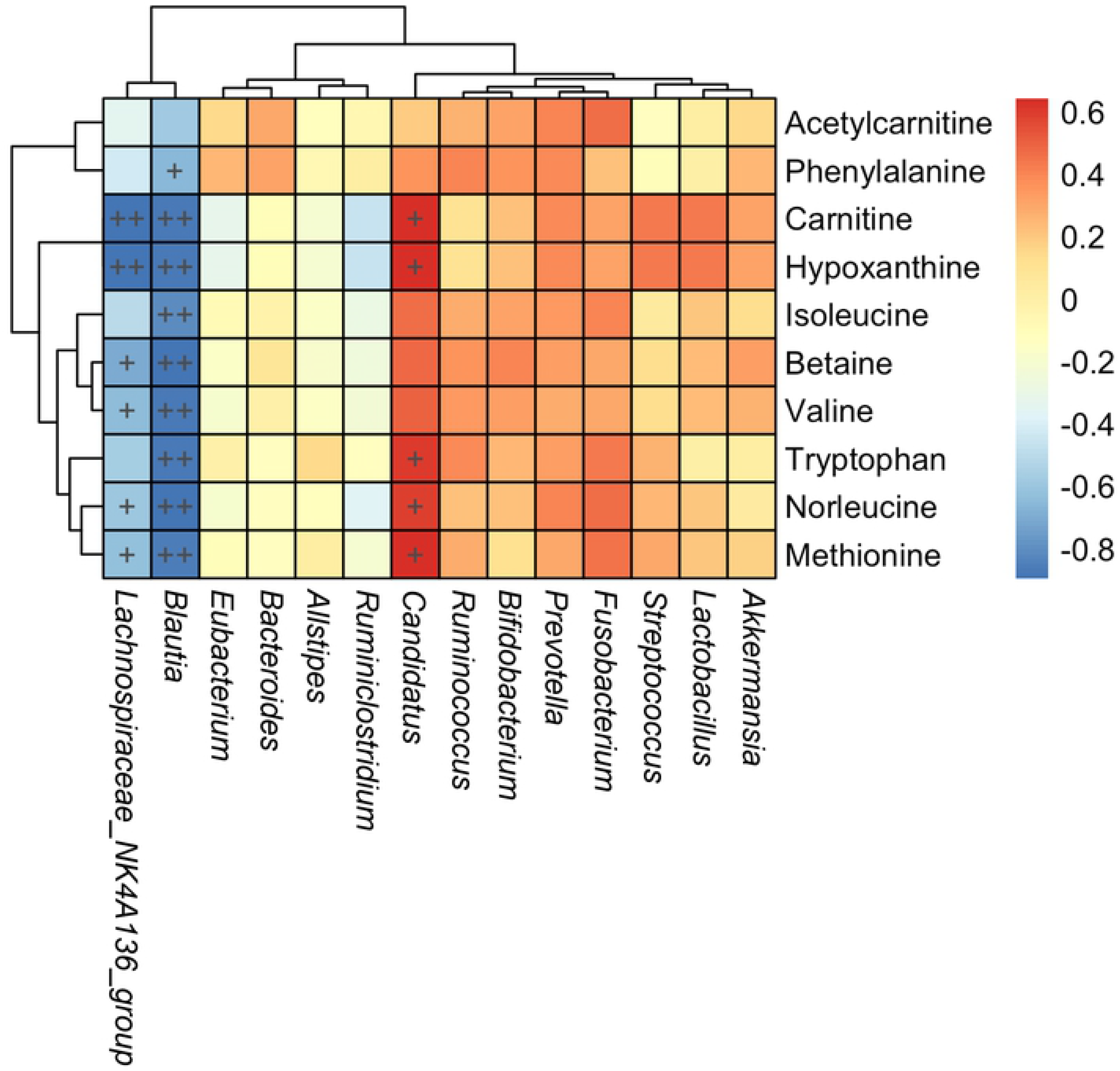
Heat map of correlations between expression of metabolites and microbiota composition. Spearman’s correlation heat map showing the correlation between significantly expressed gut metabolites, genera of gut microbiota, and selected serum metabolites. Color intensity represents the magnitude of correlation. Red represents positive correlations and blue negative correlations. + *p* < 0.05; ++ *p* < 0.01.

## Discussion

Lin et al. previously found that YGW could ameliorate airway inflammation and airway hypersensitivity in *Der p*-induced chronic allergic asthma in mice [10]. However, the action mechanism remained to be further elucidated. In this study, serum metabolomics analysis showed *Der p* markedly elevated the level of 10 metabolites — acetylcarnitine, carnitine, hypoxanthine, tryptophan, phenylalanine, norleucine, isoleucine, betaine, methionine, and valine. These metabolites are mainly related to BCAA metabolism, AAA biosynthesis, and phenylalanine metabolism. YGW administration reversed 7 of the 10 metabolites, chiefly BCAA metabolites. 16S DNA sequencing revealed that YGW profoundly changed the composition of *Der p*-induced gut microbiota, and multiple correlation analysis indicated the 10 selected metabolites and gut microbiota composition with good correlation.

Metabolomics is a comprehensive characterization of metabolites in biological systems that generates unique chemical fingerprints for specific cellular processes. In particular, metabolomics is increasingly used to diagnose disease, understand disease mechanisms, identify novel drug targets, customize drug treatments and monitor therapeutic outcomes [37]. Asthma and airway inflammation are complex and respond to infectious stimuli, which may result in a broad spectrum of possible metabolic products. Metabolomics could provide unique and novel insights into asthma. In this study, *Der p-*induced a mouse metabolic profile with 10 significantly raised metabolic signatures, including acetylcarnitine, carnitine, hypoxanthine, tryptophan, phenylalanine, norleucine, isoleucine, betaine, methionine, and valine, identified as potential biomarkers for *Der p-*induced allergic asthma. Tryptophan, phenylalanine, isoleucine, methionine, and valine are belonging to essential amino acids, which could not be synthesized by mammalian cells. Researchers found gut microbiota, such as the genera of *Bacteroides*, *Clostridium*, *Candidatus*, *Propionibacterium*, *Fusobacterium*, *Streptococcus*, *Lactobacillus*, *Akkermansia*, and *Bifidobacterium*, are contributing the source of elevated serum methionine, BCAAs (isoleucine, leucine and valine) and AAAs (tryptophan, phenylalanine and tyrosine) [38, 39]. Intestinal microbes can provide amino acids to meet host requirements, contributing to the energy delivery and modulating amino acid homeostasis [40]. A recent research also showed that HDM could cause immune system disorder with a correlation of lower diversity of gut microbiota [41]. In addition, current evidence suggested that the gut microbiota, such as *Staphylococcus*, *Rumminococcaceae*, *Lachnospiraceae_NK4A136_group*, and *Streptococcus*, have been implicated in predisposing to allergy, airway inflammation or asthma in both experimental models and clinical studies [42]. A reduction in *Bacteroidetes* and *Bifidobacteria* has been associated with the asthma [43]. Our study exhibited that some microbiota, including *Lachnospiraceae_NK4A136_group*, *Bacteroidetes, Blautia*, *Desuflovibrio* etc, were markedly lower, and some are higher in *Der p*-induced allergic asthma. After YGW treatment, the intestinal flora structure of the YGW treatment groups tended to be similar to that of the control group. Besides, the correlation analysis between metabolites and gut microbiota also demonstrated that some gut microbiota has positive correlation with amino acids changed in *Der p*-induced mouse model of chronic allergic asthma. This study provides a support that gut microbiota has a correlation with *Der p*-induced chronic allergic asthma.

Moreover, higher carnitine and phenylalanine were detected in asthma patients [44, 45]. The level of acetylcarnitine and valine were also significantly increased in ovalbumin (OVA)-induced asthma model [46], which are consistent with our results. Additionally, Ho et al. have reported recently higher tryptophan level was found in HDM-induced allergic asthma [47]. However, norleucine, isoleucine, betaine and methionine found in this study have never been previously reported to be correlated with asthma.

Cytokines/chemokines promote immune cell recruitment and activation playing an integral role in the airway inflammation. Many altered metabolites share manifold statistical associations to inflammatory cells and cytokines, implicating that they may be biologically relevant metabolic changes linked to airway inflammation. Such as hypoxanthine correlated with multiple inflammatory markers including neutrophil counts and the cytokines IL-4, IL-5, IL-6, IL-8, TNF-α, and IL-1β [48]. The present investigation found *Der p*-induced metabolic profile changes, which involved in BCAAs metabolism, AAA biosynthesis, and phenylalanine metabolism. Rising TNF-α generates inflammatory responses, induces lipolysis and increases phenylalanine fluxes [49]. Likewise, some pathogen could up-regulate interferons, which could up-regulate tryptophanyl tRNA (tRNA^Trp^) synthetase and involved in tryptophan catabolism [50]. Further, BCAAs can affect protein synthesis and decomposition, and promote glutamine synthesis. BCAAs mediate proteins, DNA, and RNA syntheses in lymphocytes and regulate immue functions [51, 52].

Malkawi et al showed that Dex could induce perturbation in several pathways, such as amino acid metabolism, pyrimidine metabolism, and nitrogen metabolism. Furthermore, Dex treatment causes significant elevation in serum levels of tyrosine and hydroxyproline, as well as significant reduction in phenylalanine, lysine and arginine [53]. Dex treatment could change the diversity and relative abundance of intestinal flora in OVA-induced asthma rats [54]. Accumulating evidences indicate that YGW is capable of modulating the immune disorders, especially boosting the immune function to strengthen the bodyline of defense against pathogen [6]. Previous study showed that YGW could attenuate *Der p*-induced inflammation via down-regulating TGF-β, IL-4, IL-5, IL-13 and inhibition of NF-κB activation [10]. In this study, we further demonstrated that YGW could improve *Der p*-induced gut dysbiosis and significantly reverse 7 raising metabolites, acetylcarnitine, tryptophan, norleucine, isoleucine, betaine, methionine, and valine, and chiefly influence valine, leucine and isoleucine biosynthesis. These results were similar to Dex. Moreover, top 10 high impact metabolites also showed a good correlation with microbial community profiling. Combined these evidences, YGW apparently ameliorates *Der p*-induced allergic asthma.

## Conclusion

In conclusion, 10 identified metabolites, acetylcarnitine, carnitine, hypoxanthine, tryptophan, phenylalanine, norleucine, isoleucine, betaine, methionine, and valine, as potential biomarkers were markedly elevated by *Der p*. Seven of the metabolites — acetylcarnitine, tryptophan, norleucine, isoleucine, betaine, methionine, and valine — could be reversed by YGW. In addition, YGW could also improve the metabolism pathway and the imbalance in intestinal flora in *Der p*-induced allergic asthma in mice. Our study provides scientific evidence for YGW administration with potential benefits for allergic asthma by ameliorating gut dysbiosis and improving the metabolome imbalance.

## Abbreviations

AA: amino acid
AAA: aromatic amino acid
BCAA: branch-chain amino acid
COPD: Chronic obstructive pulmonary disease
*Der p*: *Dermatophagoides pteronyssinus*
*Der f*: *Dermatophagoides farina*
dex: dexamethasone
HDM: house dust mites
IPA: Ingenuity Pathway Analysis
LDA: linear discriminant analysis
LEfSe: linear discriminant analysis effect size
OTUs: operational taxonomic units
PLS-DA: partial least-squares discriminant analysis
PCA: principal component analysis
PANTHER: Protein Analysis Through Evolutionary Relationships Classification System
QIIME2: Quantitative Insights Into Microbial Ecology v2
TCM: traditional Chinese medicine
VIP: variables important for the projection
YGW: You-Gui-Wan

## Declarations

### Consent for publication

Not applicable

### Availability of supporting data

All supporting data have been shown in the manuscript.

### Author contribution statement

WH Hsu: conceptualization, methodology, investigation, data curation, writing-original draft. LJ Lin and ST Kao: animal model and serum and fecal collections. YM Chao: metabolomics analysis. CK Lu: data curation. YL Lin: conceptualization, supervision, writing-revising and editing.

## Acknowledgements

This work was supported by Ministry of Science and Technology of Taiwan (MOST 106-2320-B-039-053, MOST 107-2320-B-039-001, and MOST 107-2811-B-039-519). We are grateful for Dr. Yu-Lun Kuo, Biotools Microbiome Research Center of BioTools Co., Ltd. providing assists in gut micobiota analysis.

---

**S1 Fig. High-performance liquid chromatography (HPLC) chromatogram of YGW extract**.

HPLC was run on a Hitachi 5160; 5430 DAD with 5310 oven (40°C); column: COSMOSIL 5C18-AR-II (250 x 4.6 mm); solvents: A, acetonitrile; B, 0.01% H_3_PO_3_; mobile phase: 0-15 min, 5-12% A; 15-30 min, 12% A; 30-40 min, 12-40% A; 40-45 min, 40-80% A; 45-46 min, 100 % A; detector: UV: 220 nm; flow rate: 1 ml/min.

**S2 Fig. PCA and PLS-DA analysis of metabolites with changed expression among control, *Der p* and *Der p* + YGW (0.5 g/kg) groups**

Scatter plots of scores of (A) PCA and (B) PLS-DA by LC-QTOF-MS of serum from control (blue), *Der p* (red), *Der p* + YGW (0.5 g/kg) groups (green).

**S3 Fig. Effect of YGW on TNF-α level in mouse serum**

Data are mean ± SEM. *** *P* < 0.001, \**P* < 0.05 (vs control); #*P* < 0.01, #*P* < 0.05 (vs *Der p*).

**1**: control, **2**: *Der p*, **3**: *Der p* + dex (1 mg/kg), **4**: *Der p* + YGW (0.2 g/kg), **5**: *Der p* + YGW (0.5 g/kg)

**S4 Fig. Relative abundance of the top 15 gut microbiota genera in control, *Der p* and *Der p* + YGW (0.5 g/kg) groups.**

Differential expression of genera of gut microbiota. Shows relative abundance of *Lactobacillus*, *Lachnospiraceae_NK4A136_group*, *Ruminococcus*, *Alistipes*, *Akkermansia*, *Eubacterium*, *Blautia*, *Ruminiclostridium*, *Bacteroides*, *Candidatus*, *Streptococcus*, *Staphylococcus*, *Desulfovibrio*, *Bifidobacterium*, and *Fusobacterium*. Data are mean ± SEM. \**P* < 0.05 (vs control); #*P* < 0.05 (vs *Der p*)

